# Individual differences in spatial working memory strategies differentially reflected in the engagement of control and default brain networks

**DOI:** 10.1101/2023.07.07.548112

**Authors:** Nina Purg Suljič, Aleksij Kraljič, Masih Rahmati, Youngsun T. Cho, Anka Slana Ozimič, John D. Murray, Alan Anticevic, Grega Repovš

## Abstract

Spatial locations can be encoded and maintained in working memory using different representations and strategies. Fine-grained representations provide detailed stimulus information, but are cognitively demanding and prone to inexactness. The uncertainty in fine-grained representations can be compensated by the use of coarse, but robust categorical representations. In this study, we employed an individual differences approach to identify brain activity correlates of the use of fine-grained and categorical representations in spatial working memory. We combined data from six fMRI studies, resulting in a sample of 155 (77 women, 25 ± 5 years) healthy participants performing a spatial working memory task. Our results showed that individual differences in the use of spatial representations in working memory were associated with distinct patterns of brain activity. Higher precision of fine-grained representations was related to greater engagement of attentional and control brain systems throughout the task trial, and the stronger deactivation of the default network at the time of stimulus encoding. In contrast, the use of categorical representations was associated with lower default network activity during encoding and higher frontoparietal network activation during maintenance. These results may indicate a greater need for attentional resources and protection against interference for fine-grained compared to categorical representations.

## Introduction

Research on working memory has shown that individuals use a variety of different representations and strategies to encode and maintain information over short periods of time in support of an ongoing task (e.g., Curtis, 2004; Oblak et al., 2024, 2022; Purg et al., 2022; Slana Ozimič et al., 2023; Starc et al., 2017). While mental representations describe the content of information encoded in working memory, cognitive strategies refer to the selection of mental representations and processes that are either explicitly or implicitly used by an individual to perform a working memory task (Miller et al., 2012; Oblak et al., 2024, 2022; Slana Ozimič et al., 2023). The specific representations and strategies used in working memory depend on several factors, such as the type of information to be retained (Oblak et al., 2022; Slana Ozimič et al., 2023), the type and predictability of a response to be generated (Curtis, 2004; Purg et al., 2022), the availability of attentional resources (Adam et al., 2015; Starc et al., 2017), and behavioral relevance (Klyszejko et al., 2014; Yoo et al., 2022). Increasingly, research also shows that even when faced with the same task requirements, individuals may use different representations and strategies to perform the task (Oblak et al., 2024, 2022; Slana Ozimič et al., 2023; Starc et al., 2017). Here, we investigate the neural correlates of individual differences in the use of working memory strategies in a multi-study, multi-site dataset of spatial working memory performance during functional magnetic resonance imaging (fMRI).

Spatial working memory enables the short-term storage of spatial information, such as the location of a stimulus. Extensive research has shown that memory for a stimulus location is affected by systematic distortions (e.g., Crawford et al., 2016; Huttenlocher et al., 2004, 1991). In particular, it has been observed that when individuals are asked to reproduce a stimulus location stored in working memory within an empty circle, they exhibit systematic shifts in their responses towards the diagonals of the four quadrants, formed by dividing the circle using the horizontal and vertical axes of symmetry (Huttenlocher et al., 2004, 1991). These systematic biases in spatial working memory performance have been suggested to reveal a hierarchical organization of spatial representations (Huttenlocher et al., 1991).

According to the category adjustment model (Huttenlocher et al., 1991, 2000), a stimulus location is encoded and maintained at two levels of representation – first, as a precise, fine-grained representation that stores the information of the actual location in memory, and second, as a categorical representation that assigns the stimulus location to one of a limited number of spatial categories (e.g., quadrants). The model predicts that the estimation of a stimulus location results from the combination of information at both levels, with the use of a categorical representation helping to compensate for the loss of precision in a fine-grained representation. Even though this process introduces a systematic bias in individual responses away from the correct position toward the prototypical location of the spatial category, it is assumed to increase the overall response accuracy by decreasing the variability of responses. At the neural level, the dynamic field theory (Schutte et al., 2003; Simmering et al., 2006) suggests that spatial boundaries, such as perceivable edges or spontaneously imposed axes of symmetry in task space, have a deflecting effect on memory-guided behavioral responses due to their lateral inhibitory effects, causing the activation produced by the target stimulus stored in working memory to drift in the opposite direction.

The degree of reliance on fine-grained and categorical coding of spatial locations has been related to variability in cognitive resources. In our previous work (Starc et al., 2017), we separately estimated the use of fine-grained and categorical representations during the performance of a spatial working memory task, while measuring pupil responses. We assumed that increased pupil dilation would reflect increased cognitive effort exerted toward the formation and maintenance of either fine-grained or categorical representations. Our results were consistent with a compensatory use of fine-grained and categorical representations within individuals, where a drop in attentional resources directed towards the formation of fine-grained representations during stimulus encoding resulted in increased reliance on categorical representations during late maintenance and response phases of the task. Additionally, we observed that individuals who showed on average worse fine-grained precision also exhibited greater overall use of categorical representations, suggesting stable individual differences in the use of specific representations and strategies.

Similarly, Crawford et al. (2016) found individual differences in fine-grained and categorical spatial coding that were correlated with individual spatial working memory capacity. Specifically, individuals with better spatial working memory capacity showed higher fine-grained memory precision and lower reliance on categorical representations. Since working memory capacity describes the limited cognitive resources that can be directed towards storage of information in working memory, either at the level of attentional allocation or representational capacities (Slana Ozimič and Repovš, 2020), these results suggest that individual differences in the use of fine-grained and categorical representations might be explained by the availability of cognitive resources with fine-grained representations requiring more resources than categorical representations.

Despite the extensive behavioral and computational characterization of fine-grained and categorical spatial coding, not much is known about the underlying neurobiological mechanisms. Spatial working memory is consistently characterized by sustained activation in frontal and parietal brain areas as measured with electrophysiological recordings in non-human primates (e.g., Chafee and Goldman-Rakic, 1998; Funahashi et al., 1989; Fuster, 1973; Fuster and Alexander, 1971; Kubota and Niki, 1971) and fMRI in humans (e.g., Brown et al., 2004; Courtney et al., 1998; Curtis, 2004; Srimal and Curtis, 2008; Zarahn et al., 1999). This activity is thought to reflect active engagement of these areas in working memory processes, however, the specific function of this activity has been more difficult to identify. Relating brain activity with behavioral performance of working memory tasks during fMRI has shown that brain activity varies with the level of response precision (Curtis, 2004; Hallenbeck et al., 2021), specific strategy use (Curtis, 2004; Purg et al., 2022), general memory load (Adam et al., 2018; Glahn et al., 2002; Leung et al., 2004; Linden et al., 2003; Proskovec et al., 2019) and behavioral prioritization (Klyszejko et al., 2014; Yoo et al., 2022).

In a previous fMRI study (Anticevic et al., 2010), we investigated the relationship between response accuracy in a visual working memory task and brain activity during the task. Our results showed that stronger deactivation in the temporo-parietal junction (TPJ) and the default network during stimulus encoding predicted higher accuracy of working memory performance. Since TPJ and the default network have been associated with stronger deactivation during increased cognitive effort and inhibition of distractors (Raichle, 2015a; Shulman et al., 2003; Todd et al., 2005), these results suggest that their suppression may be related to increased cognitive effort that is required to ensure good memory accuracy and protection from interference. However, the study used non-spatial visual stimuli and match-to-sample responses that do not allow the estimation of separate contribution of fine-grained and categorical representations to behavioral responses. Therefore, the brain systems and related mechanisms underlying fine-grained and categorical spatial coding have yet to be determined.

In the present study, we were interested in brain activity correlates of individual differences in the use of fine-grained and categorical representations in spatial working memory. Due to the hypothesized relationship between the use of these working memory representations and the level of cognitive resources required, we focused on brain systems that have been previously associated with general engagement of attention and cognitive control, specifically the cingulo-opercular, dorsal-attention, and frontoparietal networks (e.g., Barch et al., 2013; Cole et al., 2014; Ji et al., 2019; Raichle, 2015a; Smith et al., 2009). In addition, we investigated the role of the default network in the use of fine-grained and categorical representations, which has been associated with stronger inhibition during high attentional demands and the function of providing protection from distractors in working memory tasks (e.g., Barch et al., 2013; Cole et al., 2014; Ji et al., 2019; Raichle, 2015a; Smith et al., 2009). We hypothesized that a greater reliance on precise, fine-grained representations would be supported by increased activation of attentional and control brain systems, and a stronger inhibition of the default network. On the other hand, we assumed that uncertainty in fine-grained representations, such as due to a loss of precision or task interference, would be accompanied by a greater reliance on categorical representations that require fewer attentional and control resources.

To test these hypotheses, we investigated brain activity measured with fMRI during the performance of a spatial working memory task. A methodological challenge in the investigation of individual differences in brain-behavior relationships are low effect sizes that require large sample sizes to be detected (Elliott et al., 2020; Grady et al., 2021; Marek et al., 2022). To overcome this challenge, we combined six fMRI studies conducted at two different recording sites. Together, we used data from 155 (77 women, 25 ± 5 years) healthy individuals, which largely exceeded the average sample sizes of similar studies (e.g., around 25 participants, Marek et al., 2022). Based on behavioral performance on the task, we estimated the overall reliance on fine-grained and categorical representations of each participant by decomposing their contributions to task response errors. Individual use of fine-grained and categorical representations was then related to differences in levels of brain activity. Our results revealed individual differences in the use of spatial representations in working memory that were related to distinct patterns of brain activity. Ongoing engagement of attentional and control brain networks throughout the entire task trial, and stronger deactivation of the default network at the time of encoding a stimulus location were found to predict higher fine-grained precision in spatial working memory performance. In contrast, the use of a categorical representation was associated with lower default network activity in the encoding period and higher frontoparietal network activation in the delay period. These results suggest that the formation, maintenance and recall of fine-grained representations is supported by an increased allocation of attentional resources provided by attentional and control brain networks, whereas the categorical representations do not seem to impose such attentional demands and may be associated with an inability to protect the fine-grained representation from interference, resulting in higher reliance on the categorical representation when providing the response.

## Materials and Methods

### Participants

We combined data from six studies (Figure 1A). Three studies (Studies I-III; Table S1) were conducted at the University of Ljubljana, Slovenia, and three studies (Studies IV-VI; Table S1) at Yale University, USA. Between 11 and 37 participants took part in each study, for a total of 166 participants. All participants were healthy adults with no current or previous neurological, psychiatric, or substance-use disorders. Exclusion criteria also included contraindications to MR, such as the presence of metal implants or any other metal particles in the body, history of epileptic seizures, tremor or other motor disorders, and pregnancy. All participants had normal or corrected-to-normal vision. Several participants were excluded from further data analysis due to incomplete data collection (*N* = 5), failure to follow instructions (*N* = 1), poor data quality, or excessive movement during data collection (*N* = 2). We also excluded participants who deviated greatly from the group mean age (i.e., greater than 3 × *SD*) to ensure a more homogeneous sample (*N* = 2). Furthermore, we excluded an outlier in neuroimaging data (*N* = 1), as explained in detail in the section *fMRI acquisition, preprocessing and analysis*. Data from the remaining 155 (77 women, 25 ± 5 years) participants were used for further analysis. Most participants were right-handed (90.9%), while the rest of the participants were left-handed (11 participants, 7.14%) or ambidextrous (3 participants, 1.95%). All participants performed the behavioral task with their dominant hand. Detailed demographic information of the participants included in the data analysis are presented in Table S1. The studies carried out at the University of Ljubljana were approved by the Ethics Committee of the Faculty of Arts, University of Ljubljana, and the National Medical Ethics Committee, Ministry of Health of the Republic of Slovenia. The studies conducted at Yale University were approved by the Yale Institutional Review Board. Participants gave written informed consent before participating in the study. In all studies, participants had to perform a spatial working memory task while their brain activity was measured with fMRI.

**Fig. 1.**
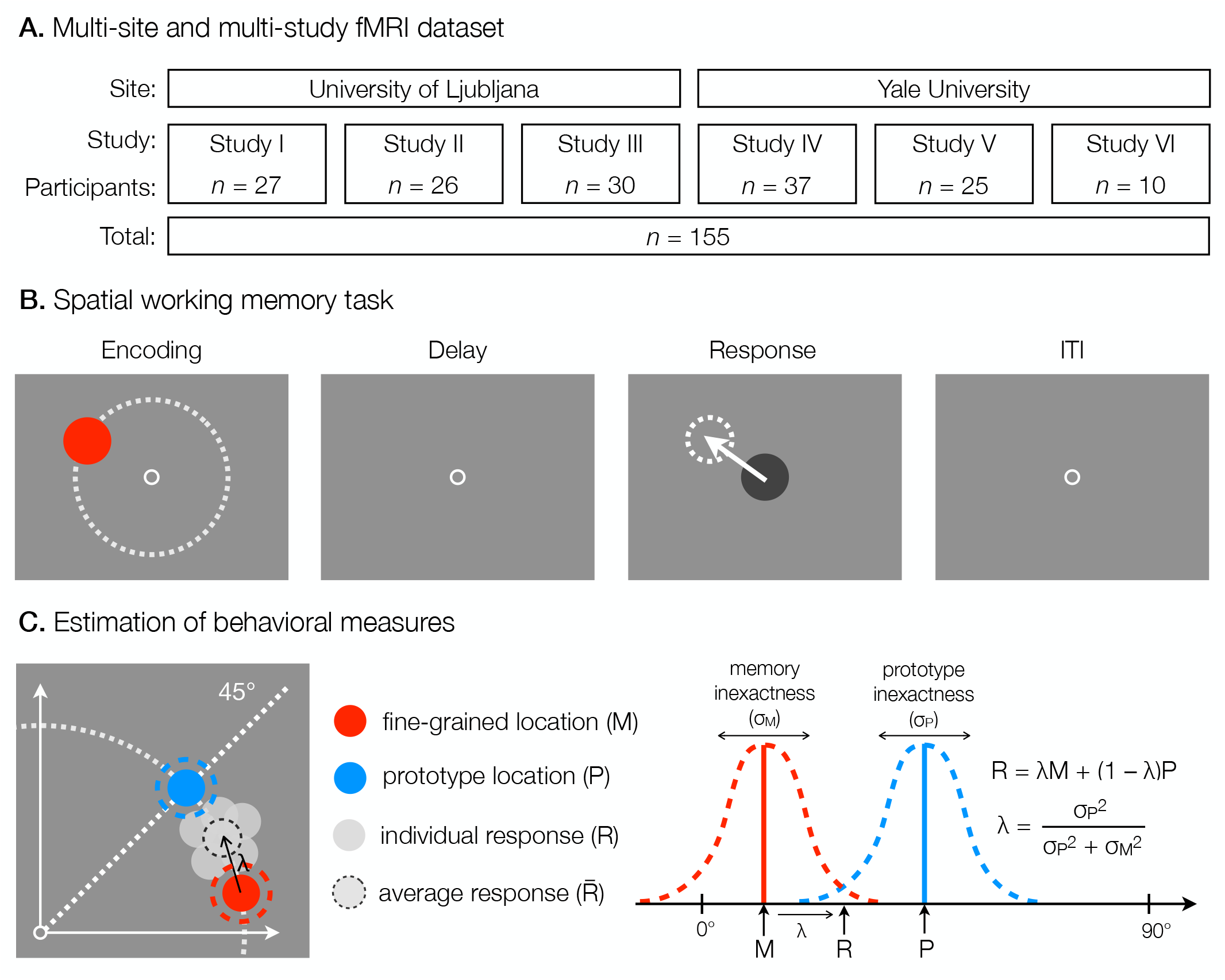
Overview of the dataset structure and behavioral methods. **A**. The dataset included six fMRI studies of spatial working memory, conducted at two different sites. In total, 155 participants were included in the data analysis. **B**. Common elements of a spatial working memory task across all studies. Each task trial consisted of a brief presentation of a target stimulus at different angles and a constant amplitude from the center of the screen, followed by a hand response to the target location using a joystick after a short delay. ITI refers to the inter-trial interval. **C**. An illustration of how the memory inexactness (*σ*_*M*_) and the prototype bias (1 −*λ*) were calculated based on angular response errors as measures of the use of fine-grained and categorical representations, respectively.

### Spatial working memory task

Individual studies were primarily conducted to address different research questions related to spatial working memory. Some of the studies are described elsewhere (Moujaes et al., 2024; Purg et al., 2022), while others are yet unpublished. The studies also differed slightly in the exact details of the spatial working memory task, which included different task conditions in each study. For the purposes of this paper, we only analyzed the task conditions that were most comparable across the studies. In particular, we focused our investigation on the task condition in which participants were asked to remember the position of a briefly presented target stimulus and, after a short delay period, to move a probe using a joystick to the position of the remembered target (Figure 1B). Despite minor differences in task design across studies, the goal of the task was always the same – a single stimulus location had to be remembered, maintained, and then recreated with a joystick on each trial. The task was displayed on an MR-compatible screen that was visible to participants from the MR scanner via a head mirror. The specific screen sizes and resolutions varied depending on the recording site and study as described in detail in Table S2. The tasks were prepared using custom scripts and run in PsychoPy (Studies I–III; Table S2; Peirce et al., 2019) or E-Prime 2.0 (Studies IV-VI; Table S2; Schneider et al., 2012). Participants responded with an MR-compatible joystick (Hybridmojo LLC, Washington, USA).

The spatial working memory task differed in the time course of task events and the exact range of target locations across studies (for details see Table S2). In three studies (Studies I-III; Table S2), the trial started with the presentation of a fixation point (2.5 s) in the center of the screen, followed by a brief presentation of a target disk stimulus. In the remaining three studies (Studies IV-VI; Table S2), the trial started immediately with the presentation of a target disk stimulus. Target stimulus presentation lasted between 0.1 s and 2 s, depending on the study. The target stimuli were presented at variable locations that were pseudorandomly selected from 20 to 36 different possible locations, depending on the study. Target locations were chosen such that the target amplitude (i.e., radial distance) from the center of the screen was constant for each participant, whereas target angles from the center of the screen varied between trials for the same participant (see Table S2 for details on the exact target amplitudes and angles for each study). The target stimuli were never presented on the cardinal axes to prevent verbalization of precise locations (Srimal and Curtis, 2008). Participants were instructed to memorize the exact position of the target stimulus. In one study (Study I; Table S2), the presentation of the target stimulus was followed by a masking pattern (0.05 s) with the aim of disrupting iconic visual memory (Curtis, 2004). In all studies, the target presentation was followed by a delay period (8 s to 10.4 s, depending on the study) during which a fixation point was presented in the center of the screen to which participants were asked to direct their gaze. In three studies (Studies IV-VI; Table S2), gaze fixation was additionally enforced by instructing participants to press a button upon a change of color of the fixation cross, which occurred randomly in 50% of trials. After the delay, a probe (i.e., a disk stimulus of the same size as the target stimulus, but a different color) appeared in the center of the screen, and participants were instructed to move the probe using a joystick to the location of the previously presented target stimulus, as precisely as possible. The time of their response was limited due to the concurrent fMRI recording between 2.3 s and 3 s, depending on the study. Individual trials were separated by an inter-trial interval (ITI) that was either fixed in duration (Studies IV-VI; Table S2) or randomly varied to allow for better task-related fMRI signal decomposition (Studies I-III; Table S2). Participants performed between 20 and 80 trials of the task, divided into 1 to 4 blocks, depending on the study.

### Behavioral data analysis

In behavioral data analysis, we first converted all behavioral data from pixel-based measurements into degrees of visual angle (°va) to provide standardization across different screen resolutions and viewing distances. At the level of individual participants, we calculated trial-to-trial response errors as the difference between the final location of the response in relation to the target location, which are thought to reflect the precision of spatial working memory. Since the findings of single-neuron recordings suggest that spatial representations are encoded at the neural level in terms of angle and amplitude in the polar coordinate system (e.g., Chafee and Goldman-Rakic, 1998; Funahashi et al., 1989; Rainer et al., 1998), we decomposed the response error on each trial into angular and amplitude differences between target and response locations measured from the center of the screen. Next, we excluded all invalid or outlier responses to ensure that the results reflected the engagement of spatial working memory and not any technical errors or inattention to the task. We defined outliers as any response that was located more than 45° away from the target location in either direction or whose amplitude was not between 0.5 and 1.75 × the target amplitude. We also excluded responses that fell outside the quadrant of the target location, defined by the horizontal and vertical axes crossing the center of the screen, to prevent the effect of misclassifying the stimulus location to the incorrect quadrant. In total, we excluded on average 2.45% of trials per participant.

During the performance of the task, only the stimulus angle was varied, while the stimulus amplitude remained constant for each participant. Thus, we assumed that memory processes would be more strongly reflected i n a ngular response errors than in amplitude response errors, and focused our further analyses on angular response errors only. To delineate the individual effects of fine-grained and categorical representations on response errors we relied on the assumptions of the category adjustment model (Crawford et al., 2016; Duffy et al., 2010; Huttenlocher et al., 2004, 1991, 2000). The model proposes that the estimation of stimulus location retained in working memory results from the combined use of fine-grained and categorical representations, each prone to decay and associated inexactness. Additionally, studies (Haun et al., 2005; Huttenlocher et al., 2004, 1991; Purg et al., 2022; Starc et al., 2017) have shown that when participants are asked to recall the position of a stimulus in a blank space, such as in the case of our study, they use four quadrants, delineated by the horizontal and vertical axes, as spatial categories, with the central value located at their corresponding diagonals, acting as the category prototype. Hence, behavioral responses collected during the spatial working memory task are assumed to be composed of a systematic shift toward the categorical center (i.e., the prototype) with the associated inexactness of this information, in addition to variability around the shifted representation due to a loss of fine-grained precision.

Computationally, we used a Bayesian model (Figure 1C), previously explained in detail in several publications (Crawford et al., 2016; Duffy et al., 2010; Huttenlocher et al., 2004, 1991, 2000), where the response (*R*) was modeled as a weighted sum of the fine-grained memory location (*M*) and the location of the categorical prototype (*P*), while the contribution of each component was defined by *λ*:

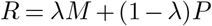

The memory location (*M*) was defined as the true target location (i.e., the target angle from the center of the screen; *µ*) with the associated standard deviation (*σ*_*M*_) reflecting memory inexactness. Similarly, the prototype location (*P*) was centered on the diagonal of the quadrant in which the target stimulus was presented (i.e., the angle of the corresponding diagonal; *ρ*), its inexactness reflected by the standard deviation around the prototype location (*σ*_*P*_). *λ* reflected confidence in the memory representation, while 1 − *λ* defined the degree of bias toward the use of prototype information. Mathematically, *λ* was defined as the ratio between the inexactness of the prototype compared to the combined inexactness of the prototype and memory representations (Crawford et al., 2016; Duffy et al., 2010; Huttenlocher et al., 1991):

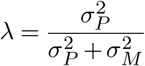

In this way, we modeled the assumption that the more inexact the memory representation is compared to the prototype, the lower the reliance on the fine-grained memory of the target and the higher the contribution of the prototype when estimating stimulus location.

The parameters of the Bayesian model were estimated using the probabilistic programming language Stan (Team, 2022b) in R (Team, 2022a). We estimated the posterior probabilities of *λ, σ*_*M*_ and *σ*_*P*_ for each participant using a two-level linear model by fitting the Student’s t-distribution to the data. Estimates were obtained based on multiple task trials per each participant, thus participants were used as a grouping variable at the first level to model varying intercepts across participants. The model was run separately for each study to prevent the potential influence of different study designs and protocols on behavioral performance. Weakly informative prior distributions were used for all model parameters, ensuring that the standard deviation of the prior distribution was at least 10 times larger than that of the posterior distribution. Specifically, we used normal prior distribution for regression parameters and half-normal distributions for standard deviations. The prior distributions were centered at mean values of the posterior parameter estimates computed with a preliminary one-level regression model to ensure stable sampling convergence. The prior distribution for the degrees of freedom parameter was set to Γ(2, 0.1) as recommended by Juárez and Steel (2010). The stability of the Hamiltonian Monte Carlo (HMC) sampling algorithm was analyzed by verifying that all estimated parameters had estimated effective sample sizes in the bulk of the distributions and in the tails of the distributions larger than 400 samples (Vehtari et al., 2021), and that the potential scale reduction statistics 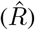 did not deviate from 1.0. To ensure stable convergence of our models, we visually inspected the trace plots of the posterior parameters and performed prior and posterior predictive checks. We verified that the maximum tree depth was not saturated. Strong degeneracies inherent to multilevel models were addressed by reparametrizing the models to a non-centered parameterization (Betancourt and Girolami, 2013).

The resulting mean estimate of *σ*_*M*_ for each participant was used as a measure of memory inexactness, since it reflected the variability around the true target location thought to result from the loss of precision in a fine-grained representation. To estimate the degree of reliance on a categorical representation, we used the measure of prototype bias defined as 1 − *λ*, which reflected the relative contribution of the prototypical location to behavioral responses, while mean *λ* was computed from posterior probabilities for individual participants.

### fMRI acquisition, preprocessing and analysis

fMRI data were collected with Philips Achieva 3TX (Studies I-III; Table S3), Siemens Tim Trio (Study IV; Table S3), and Prisma (Studies IV-VI; Table S3) scanners. We acquired T1- and T2-weighted structural images and several BOLD images using T2*-weighted echo-planar imaging sequences. We also collected pairs of spin-echo images with opposite phase encoding to estimate field maps for the purpose of distortion correction during data preprocessing. Acquisition parameters for specific images varied between different studies, as described in Table S3.

The preprocessing and analysis of the MRI data was performed with the Quantitative Neuroimaging Environment and Toolbox (QuNex; Ji et al., 2023). Several steps of analysis and visualizations were prepared using R (Team, 2022a), Matlab (R2021a, Natick, Massachusetts, USA), and Connectome Workbench (Human Connectome Project, Washington University, St. Louis, Missouri, USA).

MR images were preprocessed using Human Connectome Project (HCP) minimal preprocessing pipeline (Glasser et al., 2013). Specifically, structural images were corrected for magnetic field distortions and registered to the MNI atlas, brain tissue was segmented into white and gray matter, and the cortical surface was reconstructed. Functional BOLD images were sliced-time aligned, corrected for spatial distortions, motion-corrected, registered to the MNI atlas, and the BOLD signal was mapped to the joint surface volume representation (CIFTI) and spatially smoothed (*σ* = 4 mm). Further analyses were performed on “dense” whole-brain data (i.e., each grayordinate independently). To observe general patterns across functional brain systems and to increase statistical power, we also performed analyses on parcellated whole-brain data. Parcellated data were obtained by extracting the mean signal of 360 cortical brain regions identified based on the HCP-MMP1.0 parcellation (Glasser et al., 2016) and, additionally, for 358 subcortical regions and 12 brain networks based on the Cole-Anticevic Network Partition (Ji et al., 2019). Although the exploratory analyses were performed for all brain areas and networks, we were primarily interested in the cingulo-opercular, dorsal-attention, frontoparietal, and default networks as defined in the Cole-Anticevic Network Partition (Ji et al., 2019).

We performed the activation analysis using a general linear modeling (GLM) approach in which event regressors were convolved with the assumed double-gamma haemodynamic response function (HRF; Friston et al., 1998). For each participant, we modeled each phase of a task trial separately. Specifically, we estimated the *β* coefficients for the encoding, delay, and response phases (Figure S1). For three studies (Studies IV-VI; Table S2), we also separately modeled the attention cue in the middle of the delay period when present (Figure S1). Trials with outlier responses based on the behavioral data analysis were modeled as separate events using unassumed modeling and excluded from the group-level statistical analyses of the fMRI data. We additionally modeled motion parameters, their first derivatives, and squared motion parameters to account for any signal artifacts due to movement. To identify outlier participants based on brain activity, we computed Pearson correlation coefficients between the *β* maps for encoding, delay and response activity of each participant with a corresponding group average *β* map. We identified one participant who deviated more than 3 × *SD* from the group average *β* map and excluded this participant from further analysis.

To identify significant activation and deactivation during the task, we next analyzed the *β* estimates at the group level using permutation analysis (500 permutations, tail acceleration) in PALM (Winkler et al., 2014). To test the significance of the *β* estimates based on the “dense” grayordinate data, we conducted two-tailed one-sample t-tests with TFCE (*H* = 2, *E* = 0.5, *C* = 26) FWE correction. To test the significance of the *β* estimates based on the parcellated data, we conducted two-tailed one-sample t-tests with FDR correction. The resulting corrected *p*-value maps were thresholded at the whole-brain corrected significance level of *α <* 0.05.

### The estimation of brain-behavior relationship

To estimate the relationship between brain activity in specific networks and behavioral measures, we performed Bayesian two-level linear modeling with factors memory inexactness and prototype bias. The models were numerically estimated using the probabilistic programming language Stan (Team, 2022b) in R (Team, 2022a). To obtain standardized *β* coefficients and provide easier comparison of results across both behavioral factors, brain activity estimates, memory inexactness, and prototype bias were standardized to *µ* = 0, *σ* = 1, across all participants. We used study as the grouping variable at the first level to model varying intercepts across studies and Student’s t-distribution to describe the data. Weakly informative prior distributions were used for all model parameters, ensuring that the standard deviation of the prior distribution was at least 10 times larger than that of the posterior distribution. Specifically, we used normal prior distributions (*µ* = 0, *σ* = 10) for regression parameters and half-Cauchy prior distributions (*µ* = 0, *λ* = 2.5) for standard deviations, as recommended by Gelman (2006). The prior distribution for the degree of freedom parameter was set to Γ(2, 0.1) as recommended by Juárez and Steel (2010). The stability of the HMC sampling algorithm was analyzed by verifying that all estimated parameters had estimated effective sample sizes in the bulk of the distributions and in the tails of the distributions larger than 400 samples (Vehtari et al., 2021), and that the potential scale reduction statistics 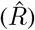 did not deviate from 1.0. To ensure a stable convergence of our models, we visually inspected the trace plots of the posterior parameters and performed prior and posterior predictive checks. We verified that the maximum tree depth was not saturated. Strong degeneracies inherent to multilevel models were addressed by reparametrizing the models to a non-centered parameterization (Betancourt and Girolami, 2013).

To examine the effect of sample size on the detection of brain-behavior relationships, we conducted Bayesian linear modeling for sample sizes ranging from 15 to 155 participants. At each sample size, 1000 samples were created by sampling with replacement from the set of all participants. We then performed Bayesian two-level normal linear model with factors memory inexactness and prototype bias with study as a random effect for each separate sample. The models were computed in the same manner as described in the previous paragraph.

## Results

### Individual differences in the use of spatial coding strategies

We first examined the pattern of response errors at different target angles in order to identify any behavioral indicators of the use of categorical representations during spatial working memory performance. We observed that participants systematically shifted their responses toward the nearest diagonals, with a greater bias occurring at target angles further away from the diagonals (Figures 2A-B and S2A). This finding indicates the use of categorical representations, where participants formed spatial categories defined by the four quadrants of the screen, delineated by the vertical and horizontal axes, each best represented by its diagonal (Huttenlocher et al., 2004, 1991; Starc et al., 2017).

**Fig. 2.**
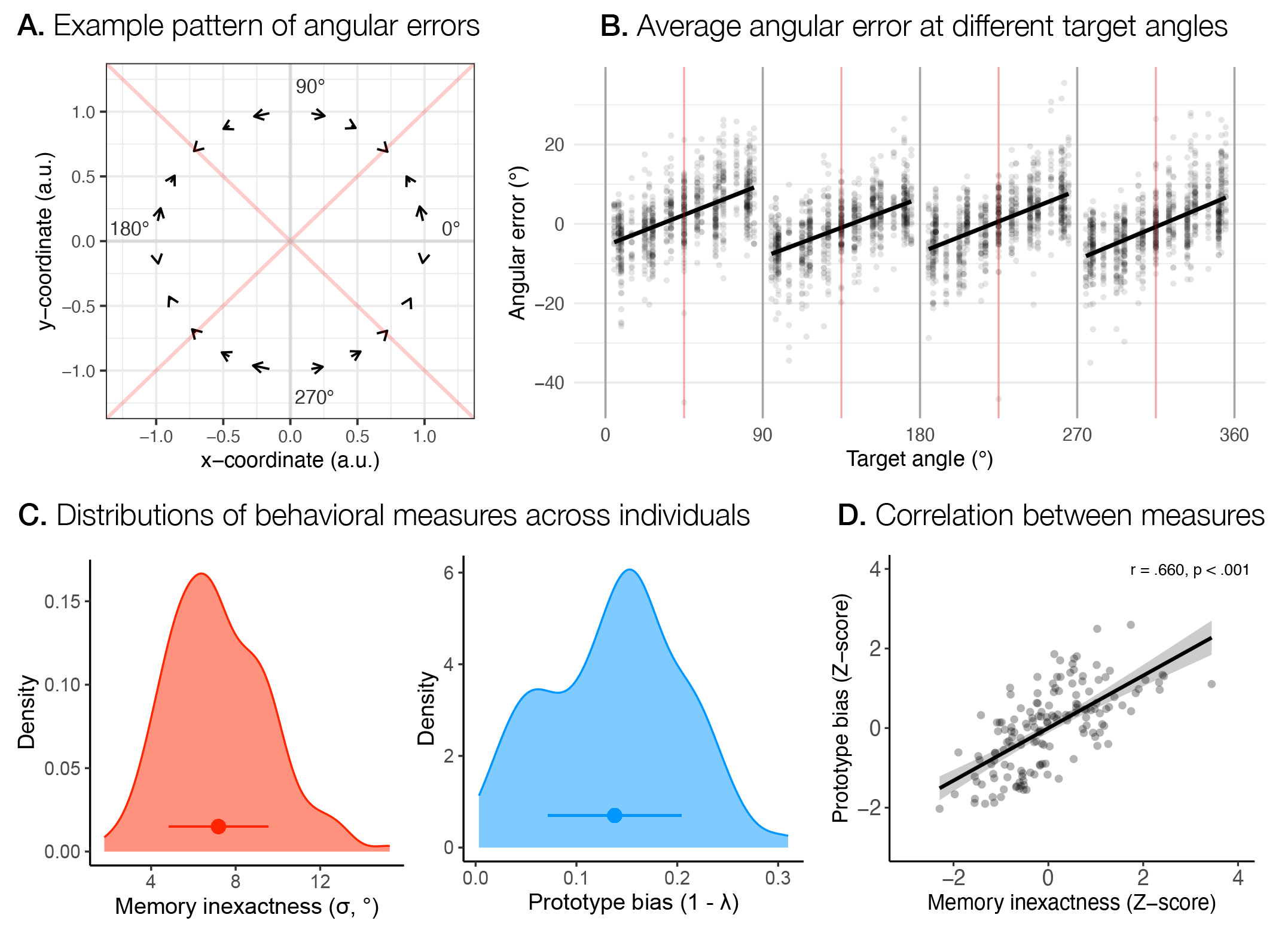
Systematic biases and individual differences in spatial working memory performance. **A**. An example pattern of the systematic bias in spatial working memory performance computed as average angular response errors at different target angles across participants in Studies IV, V and VI, which used the same stimulus angles in relation to the center of the screen. The start of the arrow denotes the target position, while the head of the arrow points to the average response position. **B**. Angular response errors at different target angles across all participants. Red lines represent diagonals of each quadrant, delineated by the horizontal and vertical axes shown as gray lines. **C**. The distribution of memory inexactness and prototype bias across participants. The points present the mean of each measure, with the range indicating the standard deviation of the measure. **D**. Relationship between memory inexactness and prototype bias across all participants estimated using Pearson correlation coefficient.

To separately measure the contribution of fine-grained and categorical representations in spatial working memory, we next derived two behavioral measures based on the modeling of the responses – memory inexactness and prototype bias. We used memory inexactness as a measure of the precision of fine-grained representations, and prototype bias as a measure of the extent to which participants relied on categorical representations. For each behavioral measure, we calculated a mean estimate for each participant, reflecting their use of fine-grained and categorical representations (for distributions across participants see Figure 2C and for study differences in both measures see Figures S2B-C). We then computed the Pearson correlation coefficient to examine the relationship between both measures across studies. Our results revealed a positive correlation, *r* = 0.660, *p <* 0.001, between memory inexactness and prototype bias (Figure 2D), suggesting that participants who relied more heavily on categorical representations showed poorer precision of fine-grained representations and vice versa.

### Task-related brain activity across different levels of parcellation

In the analysis of the fMRI data, we first examined the areas of the brain that were activated or deactivated during different phases of a task trial, namely the encoding, delay, and response phases (Figure S3A). During all phases of the trial, significant activation (i.e., *p <* 0.05 corrected for multiple comparisons) was observed in a number of brain regions, spanning the frontal, parietal, and occipital cortices. Subcortical activation was consistently observed in the cerebellum, thalamus, putamen, caudate, and brainstem. Phase-specific activations differed mainly in the early and ventral stream visual areas, where extensive activation was observed only during the encoding and response phases. Significant deactivation was observed in all phases of the trial in the posterior cingulate cortex, and in areas of the medial prefrontal cortex, and inferior frontal cortex. Additional deactivation was observed in the lateral temporal cortex for the delay and response, and in the inferior parietal cortex, early and ventral stream visual areas for the delay phase only. Subcortical deactivation was mainly observed during the delay and response phases in the cerebellum, hippocampus, and amygdala.

To investigate the integration of activity within functional brain regions and networks, and their average responses to the task, we also performed the activation analysis of the fMRI data averaged within cortical regions of the HCP-MMP1.0 parcellation (Glasser et al., 2016), and within subcortical regions and networks based on the Cole-Anticevic Network Partition (Ji et al., 2019). The results based on parcellated data showed additional significant task-related activations and deactivations (Figures 3A, S3B, and S3C). When looking at more general networks, increased activity was observed during encoding in the primary and secondary visual networks, somatomotor, cingulo-opercular, dorsal attention, frontoparietal, and language networks, in addition to the posterior and ventral multimodal networks. Deactivation was observed only in the default network. The delay phase showed significant activation in the secondary visual, somatomotor, cingulo-opercular, dorsal-attention, and posterior multimodal networks. In contrast, decreased activity was observed in the default, ventral multimodal, and orbito-affective networks during the delay. Finally, the response phase was characterized by activation in the primary and secondary visual networks, somatomotor, cingulo-opercular, dorsal attention, frontoparietal, auditory, posterior multimodal, and ventral multimodal networks. Significant deactivation was again observed only in the default network.

**Fig. 3.**
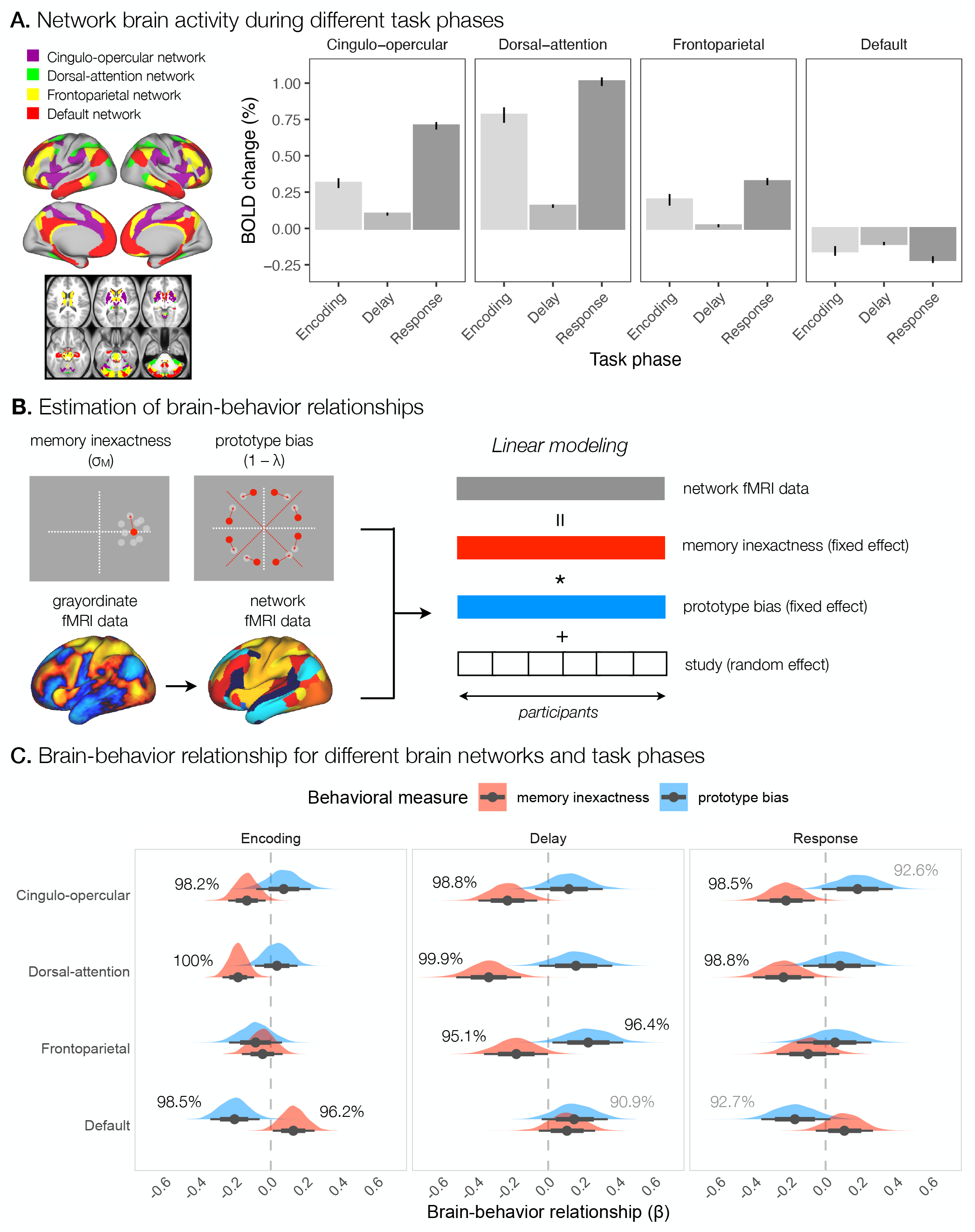
Average network activity in relation to individual spatial working memory performance. **A**. The average activity in the cingulo-opercular, frontoparietal, dorsal-attention, and default networks during different task phases. **B**. Steps in the analysis of the relationship between brain activity in specific networks and behavioral measures of memory inexactness and prototype bias. For each participant, we computed average brain activity within networks of interest defined by Cole-Anticevic Network Partition (Ji et al., 2019), and individual measures of memory inexactness and prototype bias. Next, we ran a Bayesian hierarchical linear model across participants predicting brain network activity with memory inexactness and prototype bias, and controlling for study as a random effect. **C**. Posterior distributions of the relationship between the activity of specific networks and behavioral measures of memory inexactness (red) and prototype bias (blue). Points indicate mean *β*-estimates, and lines 95% confidence intervals.

Lastly, we examined whether the analysis on parcellated fMRI data improved effect sizes or, alternatively, diluted effects due to inhomogeneous activity within individual brain regions and networks. Similar to the analysis described in Glasser et al. (2016) and Ji et al. (2019), we compared the unthresholded Z-values for delay-related activity between the “dense” grayordinate data and brain regions, and additionally between the brain regions and network data (Figure S3D). Our results showed that although the Z-values of individual grayordinates exceeded the Z-values obtained for the brain regions and networks to which they belonged, the analysis of the parcellated data resulted in higher overall effect size estimates than the analysis of the grayordinate data. Similarly, analysis of network average data resulted in higher effect size estimates than analysis of the brain regions. Although working with grayordinate data provides better spatial precision of results and is preferable when precise localization is of interest, these results suggest that working with parcellated data is preferable when testing hypotheses related to functional regions or networks, as was the case in our study.

### Individual differences in spatial coding strategies reflected in brain activity

Next, we were interested in whether individual differences in the use of fine-grained and categorical representations are reflected in brain activity. To this end, we used Bayesian linear modeling to predict the activity of brain networks of interest based on measures of memory inexactness and prototype bias (Figure 3B). Specifically, we used hierarchical linear modeling with behavioral measures as fixed factors and study as a random effect. We focused on the average activity within networks (i) to identify the engagement of broad brain systems during the use of different spatial coding strategies and (ii) to increase the effect sizes and statistical power of the analysis. Specifically, we examined brain-behavior relationships for the cingulo-opercular, dorsal-attention, frontoparietal, and default networks, separately for different task phases (Figure 3C).

During the encoding phase of the task, our results revealed 98.2% and 100% posterior probabilities for a negative relationship between memory inexactness and activity in the cingulo-opercular and dorsal-attention networks, respectively. These results suggest that increased encoding-related activity in the cingulo-opercular and dorsal-attention networks was related to decreased memory inexactness, or in other words, increased memory precision. We also observed a 96.2% posterior probability of a positive relationship between memory inexactness and activity in the default network, indicating that decreased activity in the default network was associated with increased memory precision. Relating encoding-related activity with prototype bias revealed a negative relationship between prototype bias and the default network activity with a posterior probability of 98.5%, showing that decreased activity in the default network was associated with increased prototype bias.

For the delay phase, the results indicated 98.8%, 99.9%, and 95.1% posterior probabilities for a negative relationship between memory inexactness and activity in the cingulo-opercular, dorsal-attention, and frontoparietal networks, respectively. This result again suggests that increased memory precision was related to increased activity in these networks during spatial working memory performance. On the other hand, the results showed 96.4% and 90.9% posterior probabilities of a positive relationship between prototype bias and activity in the frontoparietal and default networks, respectively. These relationships suggest that both increased frontoparietal activation and weaker deactivation of the default network are associated with increased prototype bias during the spatial working memory task.

Relating response-related activity with memory inexactness revealed a negative relationship between memory inexactness and activity in the cingulo-opercular and dorsal attention networks with posterior probabilities of 98.5% and 98.8%, respectively. The results also showed a 92.6% posterior probability of a positive relationship between prototype bias and activity in the cingulo-opercular network. These results suggest that increased response-related activity in these networks was related with increased memory precision, as well as increased prototype bias. We also observed a negative relationship between prototype bias and the default network activity with a posterior probability of 92.7%, suggesting that decreased activity in this network was associated with increased prototype bias.

The general whole-brain patterns of the relationship between brain activity and behavioral measures of memory inexactness and prototype bias for brain regions and networks can be observed in Figures S4-5. These analyses revealed several additional relationships with both behavioral measures and activity in other brain networks. For memory inexactness, a negative relationship with activity in the primary visual, secondary visual, and posterior-multimodal networks during the encoding was found with posterior probabilities of 98.9%, 100%, and 99.0%, respectively (Figure S5). We also observed a negative relationship between memory inexactness and response-related activity in the primary visual, secondary visual, somatomotor, and posterior-multimodal networks with posterior probabilities of 96.1%, 99.2%, 97.5%, and 98.7%, respectively (Figure S5). For the prototype bias, a positive relationship with activity in the language and orbito-affective networks during the delay was observed with posterior probabilities of 97.7%, and 98.8%, respectively (Figure S5).

### The effect of sample size on the detection of brain-behavior relationships

A comparatively large multi-study sample provided us with an increased power to detect brain-behavior relationships with relatively small effect sizes. To further validate the stability of the results and assess statistical power in evaluating brain-behavior relationships, we conducted a comprehensive resampling analysis. Specifically, for each sample size from 15 to 155, we randomly selected a set of participants from our original sample with replacement 1000 times and repeated the Bayesian hierarchical linear regression for the four networks of interest, i.e. the cingulo-opercular, dorsal-attention, frontoparietal, and default networks, for the delay period for each sample. This allowed us to evaluate the effects of sample size on model estimates, their confidence intervals, and statistical power.

While mean *β* coefficients estimated in the linear model were generally stable across different sample sizes (Figure S6), our results indicated that the variability of *β* estimates within each sample size changed significantly with sample size. Zero was robustly excluded from the 95% confidence interval computed across 1000 resamplings for the relationships between memory inexactness and activity in the cingulo-opercular, dorsal-attention, and frontoparietal networks only after sample sizes of 93, 73, and 151, respectively. Zero was also consistently excluded from the 95% confidence interval across 1000 resamplings for the relationship between prototype bias and the frontoparietal network activity after a sample size of 149. Statistical power, computed as the proportion of samples in which 95% of posterior distribution was above or below 0, linearly increased with increasing sample size and reached 61.3%, 87.8%, and 26.0% for the relationships between memory inexactness and activity in the cingulo-opercular, dorsal-attention, and frontoparietal networks, respectively, and 31.2% for the relationship between prototype bias and the frontoparietal network activity (Figure S6).

## Discussion

A spatial location can be encoded and maintained in working memory using different representations and strategies. Fine-grained representations provide detailed stimulus information, but are cognitively demanding and prone to inexactness. On the other hand, categorical representations may provide a more robust and less demanding strategy, but at the cost of loss of fine-grained precision. In our study, we were interested in the extent to which individuals rely on fine-grained and categorical representations to encode and maintain spatial information in working memory, and how these individual differences in spatial working memory strategies are reflected in brain activity.

### Individual differences in spatial coding strategies

The investigation of behavioral performance in the spatial working memory task revealed the presence of a systematic bias in behavioral responses. Specifically, we observed that participants tended to shift their responses closer to the nearest diagonals of the four quadrants, formed by dividing the screen at the vertical and horizontal axes of symmetry. Several previous studies (Haun et al., 2005; Huttenlocher et al., 2004, 1991; Purg et al., 2022; Starc et al., 2017) have suggested that such a bias reflects the use of categorical representations, where participants spontaneously impose spatial categories in coding stimulus position. Huttenlocher et al. (2004) have shown that this bias is replicated even when different spatial categories are imposed by the task by clustering stimuli around the horizontal and vertical axes, as well as encouraging participants to use categories centered on the cardinal axes and bounded by the diagonals. This suggests that the horizontal and vertical axes represent the most robust category boundaries, resulting in the lowest misclassification of spatial information (Huttenlocher et al., 2004). Nevertheless, the use of different reference points (Holyoak and Mah, 1982; Sadalla et al., 1980) or spatial borders (Nelson and Chaiklin, 1980; Newcombe and Liben, 1982), and specific instructions on the context of the space (Tversky and Schiano, 1989) have been shown to affect the type of categories constructed in spatial estimation tasks, suggesting that the categories formed are, at least to some extent, context-dependent (Huttenlocher et al., 1991).

Our results are in line with the category adjustment model Huttenlocher et al. (1991, 2000), which proposes that a spatial location in working memory is simultaneously represented as a fine-grained and categorical representation. The model predicts that the uncertainty in remembered fine-grained information is compensated for by using information of a broader stimulus category, which introduces a systematic bias in responses towards a prototypical value, but increases an overall response accuracy by decreasing response variability. We used the assumptions of the category adjustment model to mathematically describe behavioral responses during spatial working memory performance and to identify individual contributions of fine-grained and categorical representations to response errors. Specifically, we estimated the inexactness in fine-grained memory as a spread of responses around the true target value, while the effect of a categorical representation on the estimation of stimulus location was described in terms of a degree of systematic bias towards the prototype. We were particularly interested in the relationship between the use of both representations across individuals. Our results replicated previous observation of a positive correlation between a loss of fine-grained memory precision and the use of a categorical representation (Starc et al., 2017). Specifically, our results suggest that there are individual differences in the balance between the use of fine-grained and categorical spatial coding – individuals with higher fine-grained precision of spatial representations relied less on categorical information, whereas individuals who showed lower precision in fine-grained representations seemed to rely more strongly on categorical representations.

At the interindividual level, the degree of reliance on categorical versus fine-grained representations has been related to individual working memory capacity (Crawford et al., 2016; Stukken et al., 2016). Studies on working memory capacity have traditionally focused on estimating the number of items a participant can maintain over short periods of time by comparing task performance under different working memory loads (for a review see Luck and Vogel, 2013). However, recent studies (Bays and Husain, 2008; Spencer and Hund, 2002; Zhang and Luck, 2008) suggest that increasing the detail or precision of these objects requires additional working memory resources at the cost of reducing the number of objects that can be remembered simultaneously. Therefore, the formation of high-precision representations might be easier for individuals with a high working memory capacity, whereas a low working memory capacity would require a reduction in stimulus complexity, such as by using coarse categorical coding. Crawford et al. (2016) estimated the relationship between spatial working memory capacity and the use of fine-grained or categorical representations during spatial working memory performance based on a sample of 778 adults. Their results showed a correlation between spatial working memory capacity and different spatial coding strategies, with higher capacity predicting higher spatial precision and lower categorical bias. Moreover, consistent with these results is also the observation that introducing distractor stimuli that need to be retained during spatial working memory performance or an interference task which put additional strain on working memory resources results in an increased use of categorical representations Crawford et al. (2016); Huttenlocher et al. (1991). To sum, the use of different spatial coding strategies might be related to the availability of cognitive resources, which could explain interindividual differences in the preference for a specific strategy.

It is important to note that due to the complex hierarchical structure of our model of the effect of fine-grained and categorical representations on behavioral responses, the assumptions of the model were to some extent simplified, which could potentially affect our estimates. For example, in our model we assumed the same prototype location for all participants, which we centered on the diagonal of each quadrant. Some studies have shown that the prototype might not be located exactly on the diagonal, and might even differ between different quadrants of the task display or between participants (Huttenlocher et al., 2004, 1991). This variability in the prototype location was captured to an extent by the measure of prototype inexactness in our model, although larger incosistencies in the assumed and actual prototype location could increase the estimation of prototype inexactness and, in turn, underestimate the degree of reliance on categorical representations in spatial working memory performance.

Furthermore, our model did not account for the potential influence of inexact boundaries in the estimation of stimulus locations near boundaries. In the case of our study, participants gave their responses on a blank screen, which meant that no spatial boundary was explicitly presented, but participants spontaneously imposed boundaries in the form of horizontal and vertical symmetry axes. Their estimation of boundaries could therefore be uncertain or inexact, which could lead to misclassification of stimuli near boundaries. For instance, responses within the boundary inexactness could fall into any of the two categories delineated by the boundary and adjusted towards its center. When averaged, these responses with opposing directions of prototype bias would cancel each other out, resulting in an overall decreased effect of prototype bias around the inexact boundary (Huttenlocher et al., 2004, 1991). Our results showed that average response errors increased with target angle further from the diagonal, with a slight decrease near the boundaries, especially in the study in which the stimuli were presented the closest to the cardinal axes. For this reason, we excluded all misclassified stimuli from our data analysis to prevent dilution of the effects of categorical representations. We identified only 1.86% of misclassified stimuli per participant, with all misclassification occuring up to the target angle 15° from any boundary. The dynamic field theory (Schutte et al., 2003; Simmering et al., 2006) assumes that boundaries, perceived or spontaneously imposed, have a deflecting effect on behavioral responses during the estimation of a stimulus location in working memory due to their lateral inhibitory effects at the neural level, which results in a drift of the activation produced by the remembered target stimulus away from the boundary. Despite the overall decreased response errors near boundaries in our study, our results might still be in line with the assumptions of the dynamic field theory when looking at individual responses – i.e. the boundary might still have a deflecting effect, but in different directions for the correctly and incorrectly classified stimuli.

### Different coding strategies related to the engagement of separable brain systems

The assumed advantage of categorical spatial coding is that it is less demanding on cognitive resources without compromising the overall accuracy of responses. In contrast, encoding fine-grained information yields precise responses, but requires greater engagement of attention and cognitive control. Therefore, we hypothesized that the use of specific spatial representations would be related to the level of engagement of the attentional and control brain systems. Specifically, we expected that a stronger reliance on precise, fine-grained representations would be supported by increased activation of attentional and control brain systems, and stronger inhibition of the default network. On the other hand, we assumed that uncertainty in fine-grained representations, such as due to a loss of precision or task interference, would be related to an increased use of categorical representations that would require fewer attentional and control resources.

In the investigation of the relationship between brain activity with behavioral measures of the precision of fine-grained representations and the use of categorical representations, we observed a strong positive relationship between fine-grained memory precision and activity in the cingulo-opercular and dorsal-attention networks during all phases of the task, the encoding, delay, and response. We also observed a slightly weaker positive relationship between memory precision and the frontoparietal network activity during the delay. These results suggest that increased memory precision is indeed accompanied by an increased enagagement of these networks. The cingulo-opercular, dorsal-attention, and frontoparietal networks are consistently activated during different working memory tasks and have been widely recognized to play an important role in active maintenance of information in working memory (Brown et al., 2004; Curtis, 2006, 2004; D’Esposito and Postle, 2015; Eriksson et al., 2015; Liu et al., 2017; Purg et al., 2022; Zarahn et al., 1999). In addition, increases in the level of activity and functional connectivity within these networks have been found to scale with increased attentional demands, working memory load, and memory accuracy (e.g., Assem et al., 2020; Barch et al., 2013; Bray et al., 2015; Cole et al., 2014; Fox et al., 2005; Liu et al., 2017, 2018; Magnuson et al., 2015; Smith et al., 2009). Therefore, our findings support the notion that the formation and active maintenance of fine-grained representations presents a cognitive load and engages attentional and cognitive control systems.

Our results also revealed a negative relationship between fine-grained memory precision and the default network activity during encoding only, showing that decreased activity in this network was related to increased memory precision. Traditionally, fMRI studies investigating functional connectivity at rest have identified the role of the default network in spontaneous intrinsic activity in the absence of cognitive load (e.g., Cole et al., 2014; Damoiseaux et al., 2006; Fox et al., 2005; Greicius et al., 2003; Moussa et al., 2012; Smith et al., 2009). Moreover, the default network shows robust deactivation during the performance of various cognitive tasks, including during working memory performance, which becomes stronger with increasing cognitive load (e.g., Anticevic et al., 2010; Cole et al., 2014; Fox et al., 2005; Liu et al., 2018; Raichle, 2015a,b; Smith et al., 2009). Such decreases in the activity of the default network are thought to reflect the allocation of cognitive resources to task-relevant information and protection from distraction (Liu et al., 2017). For example, Anticevic et al. (2010) showed that stronger suppression of the default network during the encoding of target stimuli, prior to the presentation of distractors, predicted higher response accuracy in a working memory task. These results are consistent with our observations, although the study included non-spatial visual stimuli and match-to-sample responses that do not allow the uncoupling of separate contributions of fine-grained and categorical representations to response accuracy, which makes it difficult to directly relate the two studies. In summary, our results suggest that stronger inhibition of the default network is required to ensure good fine-grained memory precision, likely as a result of allocating attentional and control resources toward task-relevant stimuli and protection from interference.

Conversely, the relationship between the use of categorical representations and brain activity was somewhat less clear. Our results revealed opposing relationships with the default network activity and the prototype bias during the encoding and response phases of the task compared to the delay period. Specifically, we observed that increased use of prototype bias was related to stronger deactivation during the encoding and response, and weaker deactivation during the delay phase of the task. These results suggest temporal differences in the engagement of the default network in relation to the use of categorical representations. Stronger de-activation during the encoding and response might reflect increased attentional engagement and inhibition of distractors directed toward the formation and recall of categorical representations, respectively. On the other hand, weaker deactivation during the delay might suggest decreased attentional and control demands when individuals rely on categorical representations, supporting the hypothesis that categorical coding of spatial positions provides a less demanding spatial working memory strategy.

The investigation of the relationship between the use of categorical representations and the activity in attentional and control brain networks revealed a positive relationship between the prototype bias and activity in the frontoparietal network during the delay, as well as a slightly weaker positive relationship between the prototype bias and the activity in the cingulo-opercular network during the response. In other words, increased engagement in these networks predicted a higher use of categorical representations. While the formation, maintenance and recall of fine-grained representations required constant engagement of attentional and control systems, the results on the use of categorical representations suggest that these brain systems were engaged only later in a task trial during stimulus maintenance and recall. Similarly, Starc et al. (2017) reported a compensatory use of fine-grained and categorical representations during an individual task trial, where the failure to encode fine-grained information with high precision at the time of encoding of spatial information could then be compensated for by the reconstruction of target location based on categorical information in the late delay and response periods of the trial. These results are somewhat inconsistent with the hypothesis of reduced cognitive load and reliance on cognitive resources when using categorical representations, but may indicate a need for attentional and cognitive control during the recall of categorical representations just before the response has to be given.

However, our assumption that the observed relationship with the delay-related frontoparietal activity reflects the use of a categorical representation may be wrong. The category adjustment model (Huttenlocher et al., 1991, 2000) proposes that participants resort to the use of a categorical representation when their confidence in a memory representation is low, which would arguably be assessed just before or at the time of the response. Taking this into account, we can hypothesize that the increased frontoparietal activity does not reflect the cognitive processes engaged in the maintenance of a categorical representation, but rather the processes that predict a loss of confidence in the fine-grained memory representation and subsequent increased reliance on the categorical representation. Even though the frontoparietal network has been strongly implicated in allocation of attention and active maintenance of task-relevant information in working memory, studies have also shown its role in protection from task-irrelevant information (e.g., Jerde and Curtis, 2013; Ptak, 2012; Zhang et al., 2017). The increase in frontoparietal activity may reflect an increased effort in protecting the memory from task interference and suppression of distractors due to lower ability or confidence in the precision of the fine-grained memory representation, leading to larger reliance on categorical representation when providing the response. This is consistent with the finding that introducing an interference during the delay of spatial estimation tasks increased the reliance on categorical information (Huttenlocher et al., 2004, 1991). When assessing the role of the frontoparietal network, it is also prudent to consider the functional heterogeneity of the network. Specifically, our results of task-related activity based on voxel-wise fMRI data showed activation in some, and deactivation in other areas within the frontoparietal network, suggesting that the role of the frontoparietal network in spatial working memory processes might be more complex than initially thought, and the averaging of the activity within the network might mask diverging functions within the network.

Together, the observed patterns of associations between brain and behavior reflect important relationships between the two strategies of encoding, maintenance and recall of spatial information. While the negative relationship between fine-grained memory precision and the use of categorical representations suggests a complementary use of categorical and fine-grained representations with the goal to increase the overall response accuracy, the two strategies relate to the engagement of separable brain systems. In particular, the precision of fine-grained representations is related to the level of attentional engagement, which is reflected in the activation of the attentional and control brain networks. Additionally, greater deactivation of the default network during the formation of fine-grained representations appears to predict higher memory precision, perhaps by providing suppression of distractors and the allocation of resources toward task-relevant information. In contrast, the extent of reliance on categorical representations does not seem to impose such attentional demands. Compared with the ongoing engagement of attentional and control systems necessary to ensure high precision of fine-grained representations, some evidence was found for the activation of these systems in relation to the use of categorical representations later in the task trial during maintenance and response. Interestingly, the relationship between categorical representations and the activity in the default network appears to change over the course of the trial, where stronger inhibition of the default network is required during stimulus encoding and recall, whereas decreased inhibition is observed during the maintenance of spatial information. Since the use of a categorical representation is predicted by the uncertainty or loss of confidence in a fine-grained representation the increased deactivation of the default network during the stimulus presentation and response may reflect an increased effort in protecting the memory from task interference and suppression of distractors. On the other hand, the relaxation of the default network during the delay possibly reflects a decrease in cognitive demands in the maintenance of categorical representations.

By exploring the relationship between the use of fine-grained or categorical representations with the activity of other brain networks, we identified several relationships that suggest that the use of two strategies is related to different modalities. The precision of fine-grained memory was associated with the activity in primary visual, secondary visual, and posterior multimodal networks during the encoding and response, with increased activity in these networks predicting higher memory precision. Additionally, higher memory precision was also associated with increased somatomotor network activity during the response. These results suggest that fine-grained representations might be encoded as a visual information which is reactivated during the response, when it is converted into a motor plan used to execute the task response (Purg et al., 2022). In contrast, we observed that the use of categorical representations was predicted by the delay-related activity in the language network, with an increased categorical bias related to increased activity in this network. The engagement of the language network in the maintenance of categorical representations might indicate the transformation of spatial information into verbal codes during spatial working memory. For example, spatial categories defined as the four quadrants of the screen, delineated by the horizontal and vertical axes, could be remembered in terms of verbal codes “up-right”, “up-left”, “down-left”, and “down-right”. Similarly, studies that collected subjective reports on the strategies used during the performance of visuospatial working memory tasks have found that both, visualization and verbalization, are common strategies used to encode and maintain information in working memory (Brown and Wesley, 2013; Oblak et al., 2024, 2022; Sanfratello et al., 2014; Slana Ozimič et al., 2023). In addition, several studies have related individual differences in the use of these strategies to distinct patterns of brain activity (Kirchhoff and Buckner, 2006; Miller et al., 2012; Sanfratello et al., 2014).

In this study, we focused on general behavioral and neural strategies used in spatial working memory rather than specific mechanisms. Our results provide insight into the level of general cognitive demand involved in the use of fine-grained versus categorical representations. However, they do not indicate the specific brain regions in which the different types of information are represented. fMRI studies that used multivariate pattern analysis (MVPA) have shown that fine-grained stimulus-specific information can be decoded from early sensory areas that initially processed the stimulus (Harrison and Tong, 2009; Serences et al., 2009). In contrast, other studies have shown that the prefrontal and parietal areas can store more abstract representations, such as goals, task rules, and categories (Christophel et al., 2017; D’Esposito and Postle, 2015; Meyers et al., 2008; Riggall and Postle, 2012). Additionally, single-neuron recordings in the prefrontal cortex of monkeys during the performance of a spatial working memory task have shown that neurons, exhibiting directional selectivity for presented target angles, differed in the width of their tuning curves, suggesting that certain neurons respond to more specific directions and others to a broader range of directions (Funahashi et al., 1989). Based on these findings, it has been proposed that brain areas in the posterior-anterior axis respond to different levels of abstraction, with low-level posterior areas responding to fine-grained information and high-level anterior areas to more abstract information and regulatory signals (Christophel et al., 2017; D’Esposito and Postle, 2015; Rahmati et al., 2018). However, further studies are needed to identify areas of the brain that are involved in the storage of fine-grained and categorical representations used in spatial working memory.

### The ability to detect significant brain-behavior relationships

Several recent studies (Elliott et al., 2020; Grady et al., 2021; Marek et al., 2022; Poldrack et al., 2017) have discussed the problem of highly variable brain-behavior relationships that require large sample sizes to obtain stable and reliable results. For example, Marek et al. (2022) have shown that brain-wide association studies with typical sample sizes (i.e., around 25 participants) resulted in low statistical power, inflated effect sizes, and a failure to replicate results. We addressed this challenge by using a multi-site and multi-study fMRI dataset, which afforded us with a relatively large sample size (*n* = 155) compared to other task-related fMRI studies (Elliott et al., 2020; Marek et al., 2022). To the best of our knowledge, this is the largest fMRI dataset on spatial working memory to date. An additional advantage of a larger sample size was that it allowed us to explore the effect of the sample size on the findings of interest.

The investigation of the effect of sample size on *β* estimates as a measure of brain-behavior relationships revealed that *β* estimates can vary substantially from sample to sample when employing relatively small sample sizes. As indicated by the confidence intervals, the variability of the estimates decreased steeply at first and then slowly approached the population mean. These results are consistent with the observation that the sampling variability is large for small sample sizes and stabilizes at larger sample sizes (Marek et al., 2022). In the case of our study, brain-behavior associations appeared to stabilize roughly between 73 and 151 observations, consistent with the result obtained by Grady et al. (2021) and Schönbrodt and Perugini (2013). Moreover, similar to other fMRI studies on brain-behavior associations (Marek et al., 2022; Poldrack et al., 2017), statistical power, i.e. the ability to detect a significant effect, increased monotonically with increasing sample size, and remained fairly low even at larger sample sizes. The maximum statistical power we observed was 88.4% at *n* = 153 for the relationship between memory inexactness and activity in the dorsal-attention network.

In order to maximize sample size and statistical power, we combined data from multiple sites and studies, which presented additional challenges and limitations. Notably, there were minor differences in task designs and data collection protocols between studies, potentially contributing to the observed variability across participants. We addressed this issue using a multilevel approach. First, we analyzed the task condition that was directly comparable across studies and always had the same goal, i.e. to remember a single random target location for a few seconds on any individual trial. However, even though the task was essentially the same across studies, there could potentially be some differences in task difficulty, as a result of a different number of possible target locations, different duration of target stimulus presentation, or different length of the delay period. While we observed some differences in behavioral performance across studies, any differences in task difficulty were hard to delineate from the effects of different strategies on responses. Second, we used a hierarchical model with study as a random effect to account for any variability due to systematic differences between studies. Nevertheless, the final sample size was still relatively small compared to the recommendation of recent studies (Elliott et al., 2020; Marek et al., 2022) indicating that thousands of participants are required to prevent the inflation of effect sizes and replication failure in brain-behavior association analyses. In addition, the hierarchical structure of our data increased the complexity of the linear model used and might require even larger sample sizes to obtain reliable estimates (Kerkhoff and Nussbeck, 2019; Maas and Hox, 2004, 2005). To further increase statistical power we performed analyses on brain networks rather than grayordinates with should provide a better signal-to-noise ratio due to averaging data and ensure fewer statistical comparisons. We also used Bayesian statistical methods which have been found to give more robust results even at low sample sizes (e.g., Van De Schoot et al., 2021). Additionally, we have provided detailed power analysis to allow better insight into the stability of brain-behavior relationships in our study. However, more data or replication on an independent dataset would be welcome to further ensure the validity and generalizability of the relationships observed in our analyses.

## Conclusion

In this multi-site, multi-study analysis, we found that individuals differ in the extent to which they rely on fine-grained versus categorical representations to encode and maintain a spatial location in working memory, and that these differences correlate with the engagement of brain networks during the encoding, delay, and response phases of the task trial. Behaviorally, individuals with lower fine-grained precision relied more on categorical representations, which led to a higher categorical bias. Increased activation of attentional and control brain networks throughout the entire task trial, and stronger deactivation of the default network in the encoding period were found to predict higher precision in spatial working memory performance, possibly reflecting the importance of attentional resources for successful encoding and maintenance of the fine-grained representation. In contrast, the use of a categorical representation was associated with lower default network activity in the encoding period and higher frontoparietal network engagement in the delay period, the latter possibly reflecting an inability to protect the fine-grained representation from interference, which led to higher reliance on the categorical representation when providing the response. The results stress the need to consider individual differences in the use of specific representations and strategies when studying complex cognitive functions, such as working memory. They also illustrate the insights that the individual differences approach can provide in the study of brain-behavior relationships when a sufficient number of participants is ensured.

## Author Contributions

Conceptualization: N.P.S., J.D.M., A.A., and G.R.; Project administration: N.P.S.; Data curation: N.P.S., Y.T.C., and A.S.O.; Methodology: N.P.S., A.K., M.R., J.D.M., A.A., and G.R.; Formal analysis: N.P.S., A.K., and G.R.; Visualization: N.P.S.; Writing – original draft: N.P.S.; Writing – review & editing: N.P.S., A.K., M.R., Y.T.C., A.S.O., J.D.M., A.A., and G.R.; Supervision: J.D.M., A.A., and G.R.; Funding acquisition: A.A. and G.R.; Resources: A.A. and G.R.

## Acknowledgments

The authors would like to thank colleagues and students who helped with data collection, as well as all participants in the studies for their time and cooperation. We also thank the reviewers and editors for their constructive comments and suggestions.

## Supplementary Material

Supplementary tables and figures are available at [the link to the supplementary material].

## Data Availability

Data and analysis scripts for this paper can be found in the Open Science Framework (OSF) repository available at https://osf.io/k8mvb/.

