## Supplementary material for "Individual differences in spatial working memory strategies differentially reflected in the engagement of control and default brain networks"

### Supplementary tables

**Table S1.** Demographic data of participants included in the data analysis

| Experiment | Number | | Age (years) | | | | Handedness | | | Education (years) | | |
|---|---|---|---|---|---|---|---|---|---|---|---|---|
| | All | Females | Range | All | Females | Males | Right | Left | Both | All | Females | Males |
| I | 27 | 18 | 18–38[2] | 23.3 (5.8)[2] | 20.9 (3.0)[2] | 27.6 (7.2) | 25 | 2 | 0 | 14.5 (2.3)[4] | 13.9 (2.2)[3] | 15.8 (2.1)[1] |
| II | 26 | 17 | 19–31 | 23.0 (3.0) | 23.1 (3.5) | 22.9 (2.0) | 24 | 0 | 2 | 15.4 (2.2)[2] | 15.3 (2.6)[2] | 15.6 (1.7) |
| III | 30 | 22 | 19–42 | 24.5 (5.6) | 22.9 (4.7) | 28.8 (5.7) | 29 | 0 | 1 | 15.1 (2.7) | 14.5 (2.7) | 16.5 (2.1) |
| IV | 37 | 9 | 21–36 | 25.5 (3.4) | 25.6 (3.3) | 25.5 (3.5) | 32 | 5 | 0 | 16.7 (1.7) | 17.1 (1.6) | 16.6 (1.7) |
| V | 25 | 10 | 20–40 | 27.8 (5.9) | 26.7 (5.8) | 28.5 (6.0) | 21[1] | 3[1] | 0[1] | 16.3 (2.5) | 16.5 (2.3) | 16.2 (2.7) |
| VI | 10 | 1 | 17–31 | 22.7 (4.4) | 18.0 (–) | 23.2 (4.3) | 9 | 1 | 0 | 13.9 (2.0) | 12.0 (–) | 14.1 (2.0) |
| All | 155 | 77 | 17–42[2] | 24.7 (5.0)[2] | 23.3 (4.5)[2] | 26.1 (5.1) | 140[1] | 11[1] | 3[1] | 15.6 (2.4)[6] | 15.1 (2.6)[5] | 16.0 (2.1)[1] |

[1] Missing information for 1 participant.
[2] Missing information for 2 participants.
[3] Missing information for 3 participants.
[4] Missing information for 4 participants.
[5] Missing information for 5 participants.
[6] Missing information for 6 participants.

**Table S2.** Task parameters used in different studies

| Study | | I | II | III | IV | V | VI |
|---|---|---|---|---|---|---|---|
| Task | Trials | 36 | 32 | 24 | 20 | 20 | 80 |
| | Blocks | 2 | 2 | 3 | 2 | 1 | 4 |
| Stimuli | Diameter (px / °va) | 100 / 1.06 | 200 / 2.12 | 200 / 2.12, 2.83 | 125 / 1.72 | 125 / 1.72 | 125 / 1.72 |
| | Angles (°) | 5–355 (steps of 10) | 7.5–352.5 (steps of 15) | 7.5–352.5 (steps of 15) | 9–351 (steps of 18) | 9–351 (steps of 18) | 9–351 (steps of 18) |
| | Amplitude (px / °va) | 400 / 4.24 | 400 / 4.24 | 400 / 4.24, 5.66 | 415 / 5.72 | 415, 390 / 5.72, 5.38 | 415, 390 / 5.72, 5.38 |
| Events | Fixation (s) | 2.5 | 2.5 | 2.5 | – | – | – |
| | Target (s) | 0.1 | 2 | 2 | 1.4 | 1.6 | 1.6 |
| | Mask (s) | 0.05 | – | – | – | – | – |
| | Delay (s) | 9.85 | 8 | 8 | 9.8 | 10.4 | 10.4 |
| | Attention cue (s) | – | – | – | 1.4 | 1.6 | 1.6 |
| | Response (s) | 3 | 3 | 3 | 2.8 | 3.2 | 3.2 |
| | ITI (s) | 12.5, 15, 17.5 | 12.5, 15, 17.5 | 12.5, 15, 17.5 | 13.3 | 15.2 | 15.2 |
| | ITI ratio | 3:2:1 | 5:2:1 | 5:2:1 | – | – | – |
| Screen | Size (mm) | 640 x 400 | 640 x 400 | 640 x 400 | 427 x 343 | 427 x 343 | 427 x 343 |
| | Resolution (px) | 2560 x 1600 | 2560 x 1600 | 2560 x 1600, 1920 x 1200 | 1280 x 1024 | 1280 x 1024 | 1280 x 1024 |
| | Viewing distance (mm) | 1350 | 1350 | 1350 | 1385 | 1385 | 1385 |

**Table S3.** MRI parameters used in different studies

| Study | | I | II | III | IV | V | VI |
|---|---|---|---|---|---|---|---|
| **Scanner** | | Philips Achieva 3.0T TX | | | Siemens 3T Tim Trio or Prisma | | |
| **T1w and T2w** | Sagittal slices | 236 | 236 | 236 | 224 | 208 | 208 |
| | FOV (mm) | 224 x 235 | 224 x 235 | 224 x 235 | 256 x 256 | 256 x 256 | 256 x 256 |
| | Voxel size (mm) | 0.7 | 0.7 | 0.7 | 0.8 | 0.8 | 0.8 |
| | TR (ms) | T1: 12, T2: 2500 | T1: 12, T2: 2500 | T1: 12, T2: 2500 | T1: 2400, T2: 3200 | T1: 2400, T2: 3200 | T1: 2400, T2: 3200 |
| | TE (ms) | T1: 5.7, T2: 414 | T1: 5.7, T2: 403 | T1: 5.7, T2: 403 | T1: 2.07, T2: 564 | T1: 2.22, T2: 563 | T1: 2.22, T2: 563 |
| | Flip angle (°) | T1: 8, T2: 90 | T1: 8, T2: 90 | T1: 8, T2: 90 | T1: 8, T2: T2 var | T1: 8, T2: T2 var | T1: 8, T2: T2 var |
| **BOLD** | Axial slices | 48 | 48 | 48 | 54 | 72 | 72 |
| | FOV (mm) | 240 x 240 | 240 x 240 | 240 x 240 | 210 x 210 | 208 x 208 | 208 x 208 |
| | Voxel size (mm) | 3 | 3 | 3 | 2.5 | 2 | 2 |
| | TR (ms) | 2500 | 2500 | 2500 | 700 | 800 | 800 |
| | TE (ms) | 27 | 27 | 27 | 31 | 37 | 37 |
| | Flip angle (°) | 90 | 90 | 90 | 55 | 52 | 52 |
| | SENSE factor 2 | 2 | 2 | 2 | – | – | – |
| | Multi-band factor | – | – | – | 6 | 8 | 8 |
| | Number of runs | 2 | 2 | 3 | 2 | 1 | 4 |
| | Frames per run | 215 | 189 | 281 | 400 | 770 | 770 |
| **Field maps** | Axial slices | 48 | 48 | 48 | 54 | 72 | 72 |
| | FOV (mm) | 240 x 240 | 240 x 240 | 240 x 240 | 210 x 210 | 208 x 208 | 208 x 208 |
| | Voxel size (mm) | 3 | 3 | 3 | 2.5 | 2 | 2 |
| | TR (ms) | 2500 | 2500 | 2500 | 731 | 8000 | 8000 |
| | TE (ms) | 27 | 27 | 27 | 4.92/7.38 | 66 | 66 |
| | Flip angle (°) | 90 | 90 | 90 | 50 | 90 | 90 |
| **Other physiological data** | | – | EEG | EEG | – | eyetracker | eyetracker |

### Supplementary figures

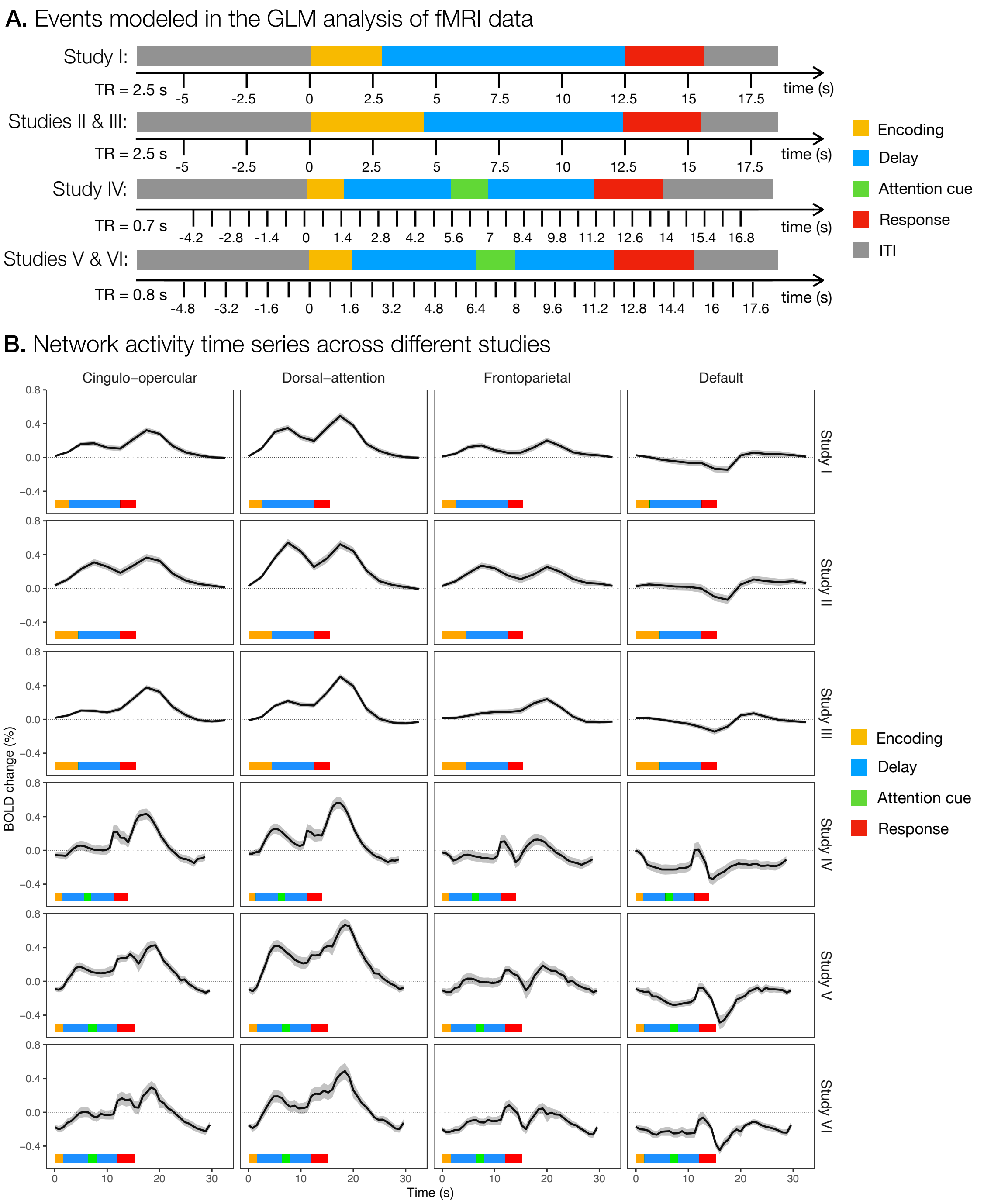


**Figure S1. Linear modeling of the brain activity during a task trial. A.** The timeline of events during a task trial modeled in the fMRI data analysis. Zeros mark the start of a task trial. **B.** The average activity is shown for the cingulo-opercular, dorsal-attention, frontoparietal, and default networks. The shaded area represents the standard error. Colored rectangles mark the timing of different events during a task trial in different studies.

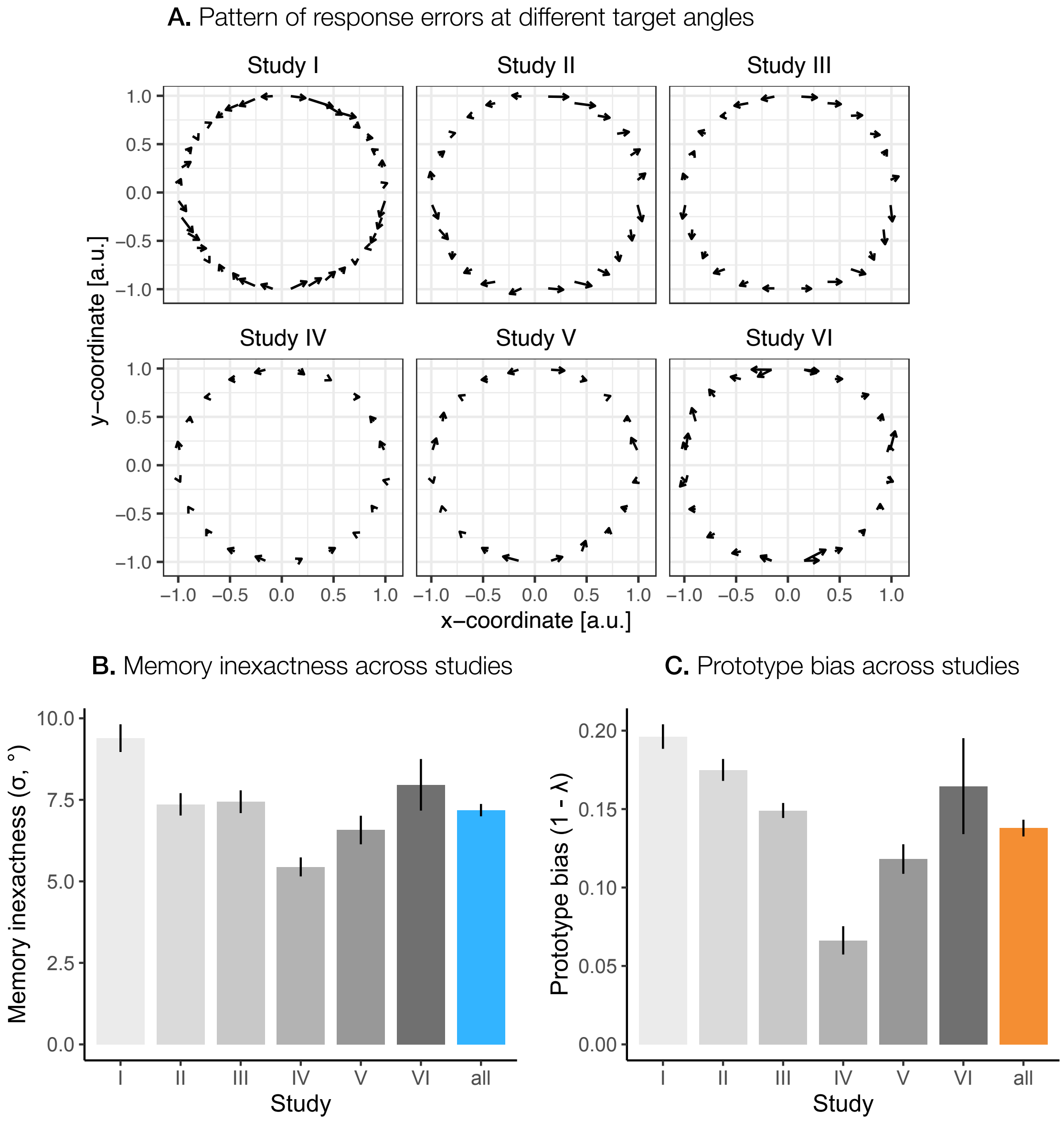


**Figure S2. Systematic biases in behavioral performance. A.** Pattern of average response errors at different target angles for individual studies. The start of the arrow denotes the target position, while the head of the arrow points to the average response position. **B.** Average memory inexactness across all participants and for individual studies. The error bars represent the standard error. **C.** Average prototype bias across all participants for individual studies. The error bars represent the standard error.

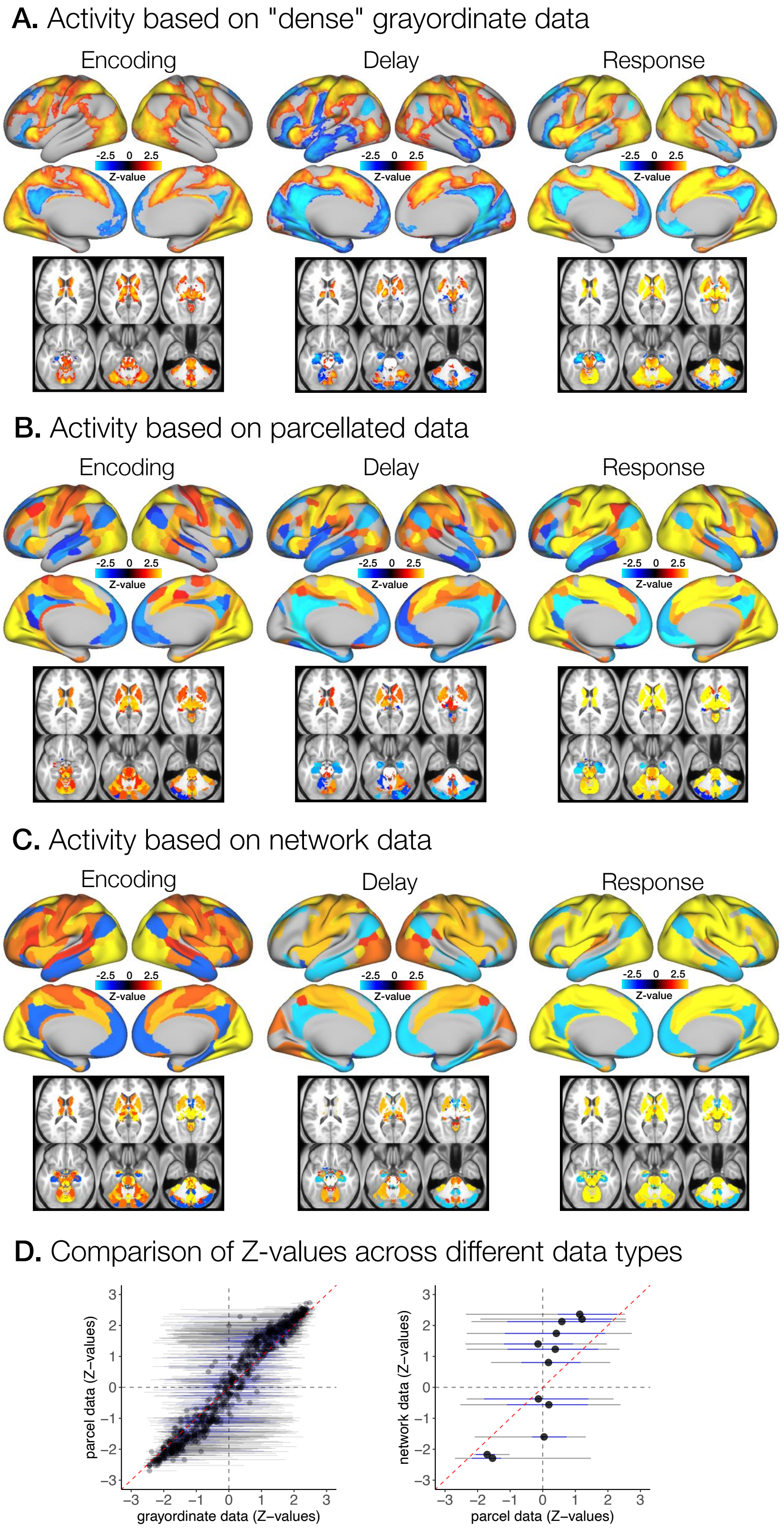


**Figure S3. Brain activity during different phases of a task trial based on different levels of fMRI data parcellation.** Significant activation and deactivation during the encoding, delay, and response phases for **A.** "dense" grayordinate, **B.** brain parcel, and **C.** network fMRI data. $p$-values for "dense" grayordinate data were corrected for multiple comparisons with TFCE FWE, whereas $p$-values for parcel and network fMRI were corrected with FDR. All images were thresholded at $p < 0.05$. **D.** The comparison of unthresholded Z-value maps for delay-related activity between "dense" grayordinate and parcel data, and additionally, for parcel and network data. The gray lines represent the range, the blue lines the inter-quartile interval (IQR), and the red dashed line the diagonal.

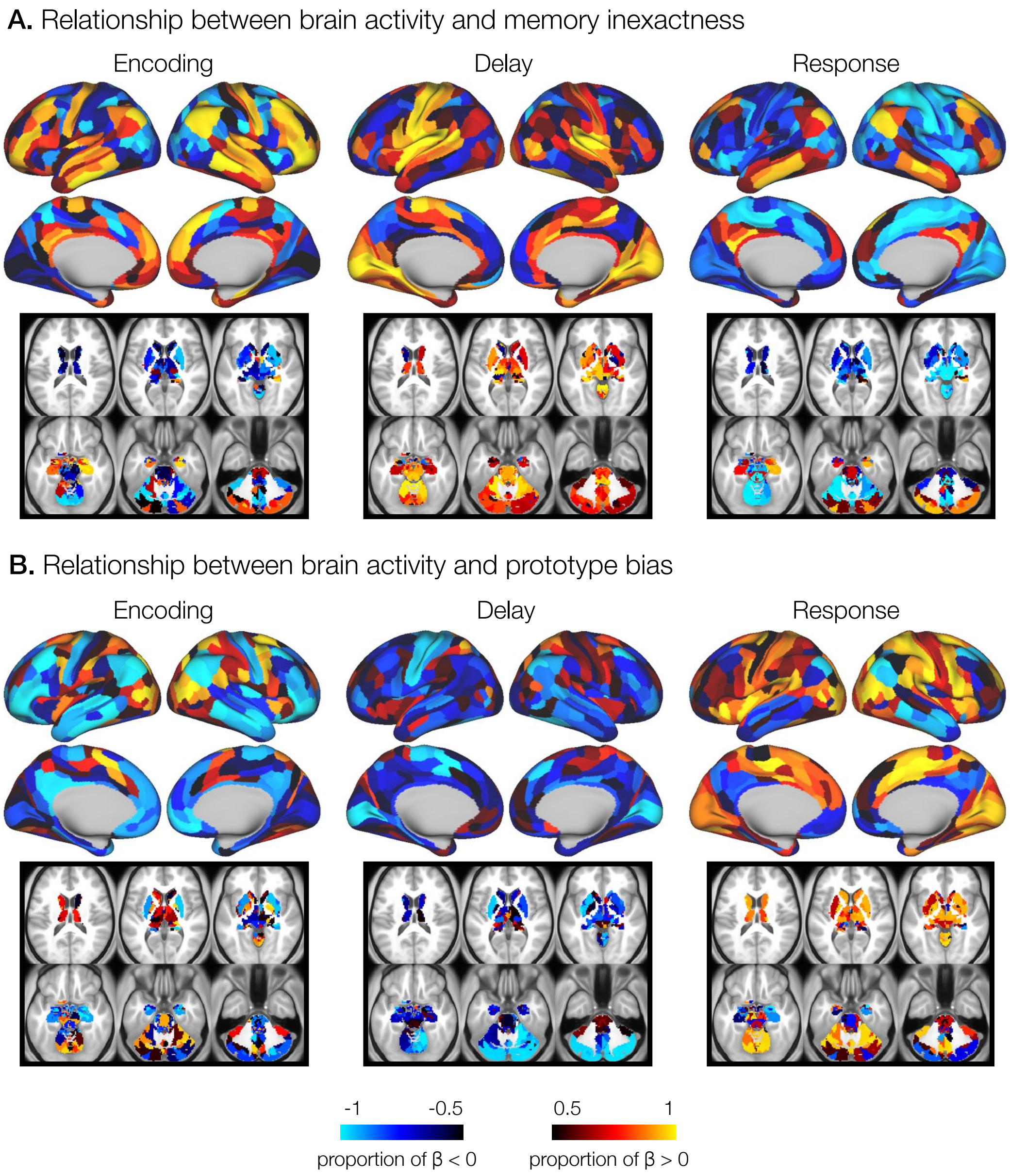


**Figure S4. The relationship between brain activity based on brain region fMRI data and individual behavioral performance.** The relationship between activity and behavioral measures was estimated by running a Bayesian linear model across participants with factors memory inexactness and prototype bias, and study as a random effect for each task phase separately. Shown are the proportions of the posterior distributions below or above $0$ for the relationship between the activity of specific networks and behavioral measures of memory inexactness and prototype bias. The results are presented for the encoding, delay, and response phases of a task trial.

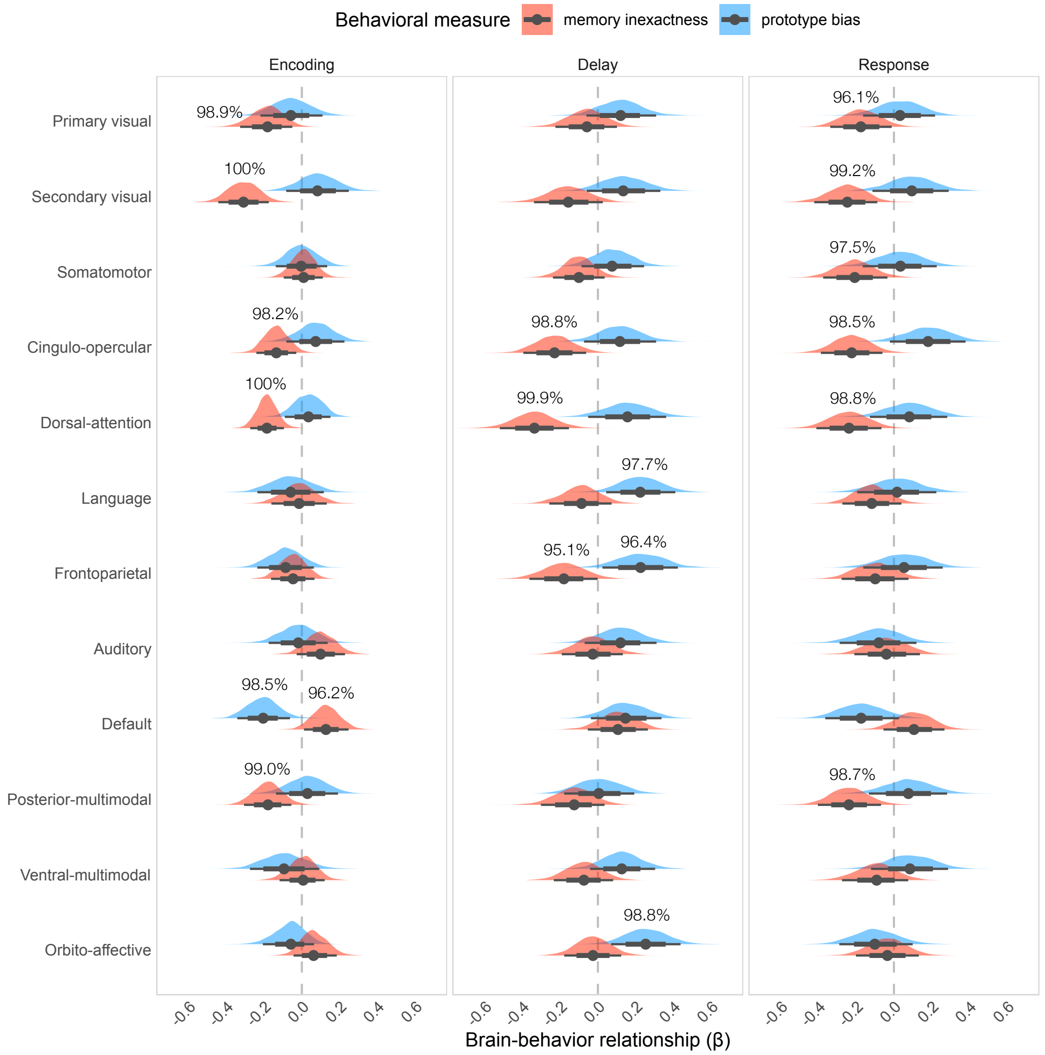


**Figure S5. The relationship between brain activity based on network fMRI data and individual behavioral performance.** The relationship between activity and behavioral measures was estimated by running a Bayesian linear model across participants with factors memory inexactness and prototype bias, and study as a random effect for each task phase separately. Shown are posterior distributions of the relationship between the activity of specific networks and behavioral measures of memory inexactness (red) and prototype bias (blue) for the encoding, delay, and response phases of a task trial. Points indicate mean $\beta$-estimates, and lines $95\%$ confidence intervals.

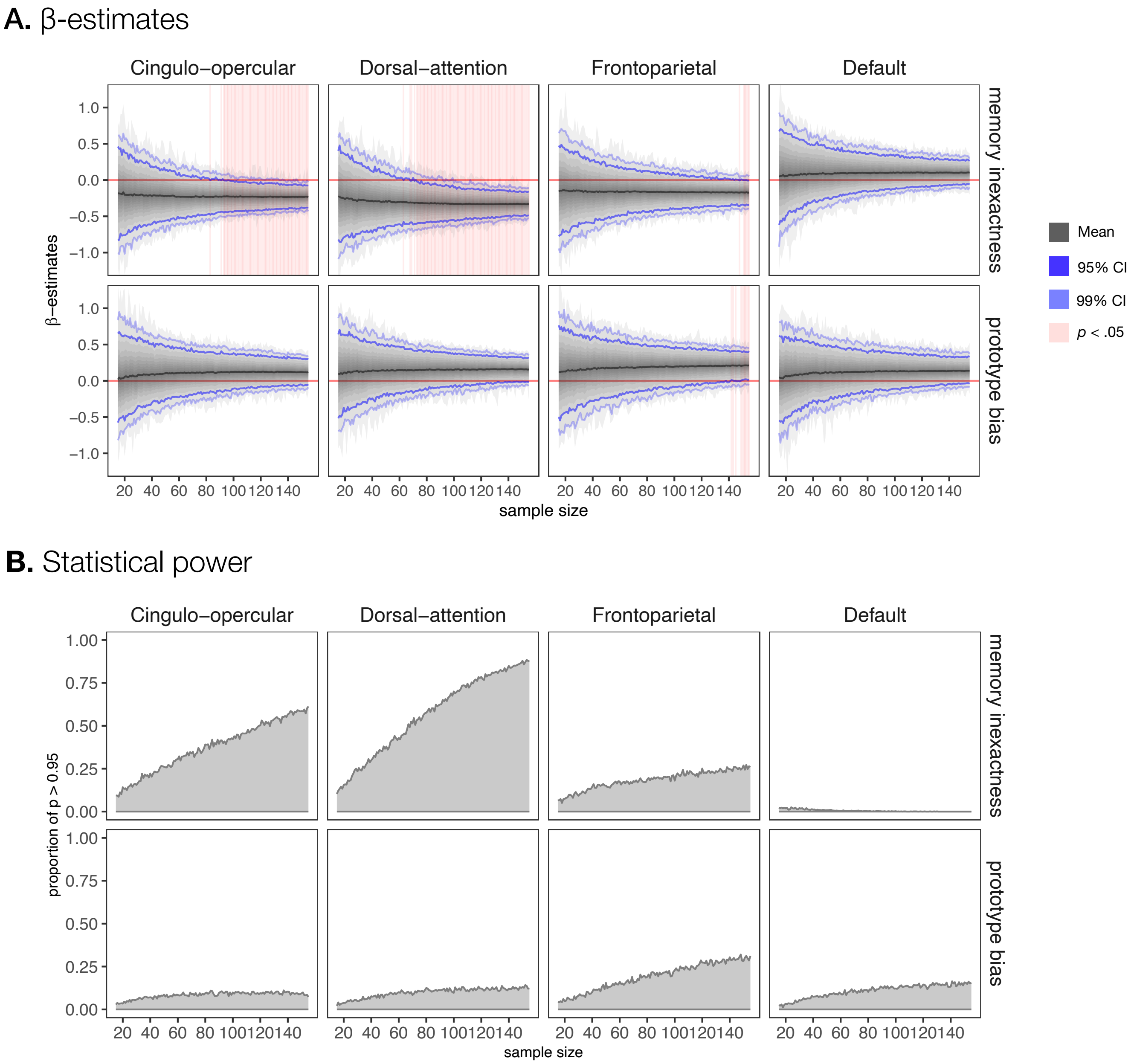


**Figure S6. The effect of sample size on the estimation of brain-behavior relationships.** We investigated the effect of sample size on **A.** $\beta$-estimates and **B.** statistical power in the investigation of the relationship of brain activity and behavioral measures of memory inexactness or prototype bias. The posterior probability of brain-behavior relationship was estimated using a Bayesian two-level normal linear model with factors memory inexactness and prototype bias, while the study was used as the grouping variable on the first level to model varying intercepts across studies, based on a varying number of sample sizes from $15$ to $155$. At each sample size, $1000$ samples were created, each by sampling with replacements from the set of all participants. **A.** The black line denotes mean across all samples, the grayed area denotes the span between maximum and minimum value with the darker shading reflecting higher density, the lighter blue line denotes the upper and lower boundary for $99\%$ of samples, the darker blue denotes the upper and lower boundary for $95\%$ of samples. The red line denotes the value $0$, the pink background shading denotes the sample sizes for which the $95\%$ confidence interval does not include $0$. **B.** The proportion of samples where $95\%$ of posterior distribution was above or below $0$.
